# Effect of systemic treatment with N-[2-(5-hydroxy-1H-indol-3-yl)ethyl]-2-oxopiperidine-3- carboxamide (HIOC) or tauroursodeoxycholic Acid (TUDCA) on retinal ganglion cell death following optic nerve crush

**DOI:** 10.1101/733568

**Authors:** Ying Li, Xian Zhang, Jiaxing Wang, Jana T. Sellers, Amber P. Boyd, John M. Nickerson, Jeffrey H. Boatright

**Affiliations:** Center for Visual and Neurocognitive Rehabilitation, Atlanta Veterans Administration Health Care System, Decatur, GA, United States; Department of Ophthalmology, Emory University School of Medicine, Atlanta, Georgia

**Author notes:** Correspondence to: Jeffrey H. Boatright, PhD, Atlanta VA Health Care System, 1670 Clairmont Road, Decatur, GA 30033 or Department of Ophthalmology, Emory University, B5500, Clinic B Building,1365B Clifton Road, NE, Atlanta, GA, 30322.

## Abstract

The aim of this study was to investigate the protective effects of systemically administered N-[2-(5-hydroxy-1H-indol-3-yl)ethyl]-2-oxopiperidine-3-carboxamide (HIOC) or tauroursodeoxycholic acid (TUDCA) in an optic nerve crush (ONC) mouse model. HIOC (50 mg/kg) or TUDCA (500mg/kg) were intraperitoneally (i.p.) injected into adult C57BL/6 mice three times per week. Two weeks after treatment (6 injections), unilateral optic nerve crush was conducted followed by treatment at the same day. The treatment was continued until 1 week or 2 weeks after ONC. A control cohort was identically treated with drug vehicle (phosphate-buffered saline; PBS). Retinas were harvested for whole mount immunofluorescence staining with RGC markers and imaged by fluorescent confocal microscopy at 40x magnification. Fluorescing cells were counted by computer-assisted automated identification and counting software (CellProfiler). Cohort sampling sizes were N=4 and statistical tesing was by the Wilcoxon-Mann-Whitney Test. Significant loss (80%’85%) occurred in the PBS-injected group 1 and 2 weeks after ONC. This loss was partially but significantly prevented in drug-treated cohorts (P < 0.05). Delivery of HIOC or TUDCA by i.p. injection increased survival of RGCs after ONC. Protection was similar between treatment with either drug. These data suggest that it is worthwhile to further explore possible protective effects of HIOC or TUDCA on RGC subtypes with regards to structure and function and in additional disease models that involve RGC loss.

## Introduction

Glaucoma is a group of diseases of the visual system that leads to the death of retinal ganglion ells and has beenestimated to affect 67 million people worldwide [1]. The most common form, primary open-angle glaucoma (POAG), accounts for approximately 12% of all cases of blindness in the United States [1]. Glaucoma is traditionally defined by elevated intraocular pressure, optic disc cupping, and/or visual field loss. However, the hallmark feature common to all types of glaucoma is the loss of retinal ganglion cells.

The optic nerve crush model developed by Schwartz et al. causes axonal injury to the optic nerve that can be evaluated morphologically and functionally [2]. This damage inflicted directly upon the optic nerve fibers leads to the degeneration and eventual death of their cell bodies by apoptosis [3, 4]. Current pharmacological therapy largely focuses on the lowering of the intraocular pressure, yet there is an increasing interest in neuroprotection. In 2001, Naskar, et al. demonstrated that phenytoin was able to rescue RGCs from secondary degeneration using the crush model in rats [5, 6]. In addition, the systematic administration of the second-generation tetracycline derivative, minocycline, has been proven to delay the death of RGCs in rats after the optic nerve damage [7, 8].

Tauroursodeoxycholic acid (TUDCA) is an endogenous bile acid that has recently shown remarkable potential to block apoptosis following cellular injury and several models of neurodegeneration and damage, including models of retinal degeneration [9–16]. TUDCA is neuroprotective in a transgenic mouse model of Huntington’s disease [17], in rat models of ischemic and hemorrhagic stroke [9, 18], and in a rat model of Parkinson’s disease (Duan et al., 2002). The mechanism of actions includes stabilizing the mitochondrial membrane, and thus inhibiting apoptosis [10], and prevention of endoplasmic reticulum (ER) stress by acting as a chaperone [16, 19, 20].

The rodent optic nerve crush model was used in the present study to induce retinal ganglion cell death. The damage that occurs mimics traumatic optic neuropathy (TON), a condition that causes a partial or complete loss of optic nerve function and, usually, immediate loss of vision. The loss of RGCs occurs in a way that is comparable to glaucoma, regarding the crush model as a highly acute glaucoma model [21]. Given that TUDCA has been proven to prevent degeneration of neuronal cells by suppressing apoptosis (Rodrigues, et al. 2000), we investigated the protective effects of TUDCA on retinal ganglion cells following optic nerve crush.

## Methods

### Animals

All animal experiments were performed in accordance the ARVO Statement for the Use of Animals in Ophthalmic and Vision Research. Adult male and female C57/BL6 mice were used for all experiments. The mice were maintained in standard housing with food and water provided ad libitum and a 12-hr light/dark cycle. Each animal was anesthetized using an 80 mg/mL ketamine hydrochloride/12 mg/mL xylazine hydrochloride solution (Sigma-Aldrich Corp, St. Louis, MO).

### TUDCA Preparation & Administration

Tauroursodeoxycholic Acid, Sodium Salt (Calbiochem, EMD Biosciences, Inc, La Jolla, CA) was dissolved in 0.15M NaHCO_3_. The pH of the solution was brought to 7.4 - 7.5 using 1M HCl or 0.15M NaHCO_3_, yielding a final concentration of approximately 50mg/mL. The mice were divided into TUDCA and vehicle (0.15M NaHCO_3_) treatment groups. Each animal received subcutaneous injections of 500mg/kg body weight of TUDCA or vehicle 24 hours prior to, the day of, and every three days following optic nerve crush.

### Optic Nerve Crush

Using a binocular operating microscope, the conjunctiva of each eye of the anesthetized mice was laterally excised. The optic nerve was visualized by subconjunctival soft dissection.. A longtitudal incision was then made along the meningial sheath, and the optic nerve crushed 2 mm posterior to the globe. Care was taken to preserve the retinal blood supply. Goniosol (CIBA Vision Ophthalmics, Atlanta, GA) was administered post-operatively as needed to relieve dryness. Buprenex (Reckitt Benckiser Pharmaceuticals Inc., Richmond, VA) and Banamine (Schering-Plough, Omaha, NE) were administered for pain and inflammation, respectively.

### Retinal Ganglion Cell Survival Analysis

Fluorogold, introduced by Schumed and Fallon in 1986, is a neuronal retrograde tracer Therfore, to visualize RGC viability and the effect of TUDCA treatment, RGCs were retrogradely labeled with fluorescent dye by placing a small piece of Surgifoam (Ethicon, Sommerville, NJ) soaked in 10 % Hydroxystilbamidine (Molecular Probes, Eugene, OR) onto the damaged optic nerve of each eye immediately following crush. Animals were sacrificed by carbon dioxide asphyxiation. Eyes were enucleated at 5, 7 & 10 days post-optic nerve crush and fixed in 10% buffered formalin (Stephens Scientific, Riverdale, NJ) for 30 minutes at 4°C in total darkness to prevent possible bleaching of signal. Following a PBS wash, the retinae were removed and flat mounted onto glass slides. The slides were cover slipped and the edges were sealed using nail polish. Images were captured using a Nikon Eclipse E800 microscope (Nikon, Melville, NY) using a DAPI filter with an excitation of 360/40. The retina was divided into three different eccentricities from the optic disc to count the viable RGCs: central (’0.5mm), middle (’2mm), and peripheral (’4mm), and separated into the superior, inferior, nasal and temporal retina. The fluorescing RGCs were then counted using ImageJ (NIH, Bethesda, MD). The data obtained from each retina were pooled for comparison across regions. The data obtained in all experiments was analyzed for mean and standard error of the mean (S.E.M.).

In other experiments, retinas were harvested for whole mount immunofluorescence staining with RGC markers Brn3a or RPBMS and imaged by fluorescent confocal microscopy at 40x magnification. Fluorescing cells were counted by computer-assisted automated identification and counting software (CellProfiler). Cohort sampling sizes were N=4 and statistical tesing was by the Wilcoxon-Mann-Whitney Test.

### Apoptosis Detection

Apoptosis in RGCs was assessed in eye sections by fluorescent TUNEL (TdT-mediated dUTP Nick-End Labeling) using the DeadEnd Fluorometric TUNEL Assay kit (Promega Corp., Madison, WI) and by immunohistochemical staining with an antibody specific for activated caspase-3 (Anti-ACTIVE Caspase-3 antibody; Promega) as previously described [Boatright et al., 2006; Chang et al., 2007]. Briefly, eyes were fixed in paraffin and cut on a microtome into 5µm sections bisecting the optic disc superiorly to inferiorly. For TUNEL, sections were deparaffinized, rehydrated, treated with Proteinase K, reacted with fluorescein-12-dUTP labeled TdT/nucleotide mix, and counter-stained using propidium iodide (Sigma-Aldrich Corp., St. Louis, MO). For active caspase-3 detection, sections were permeabilized in 0.1% Triton X-100/PBS, followed by an incubation with Anti-ACTIVE Caspase-3 antibody and fluorescein conjugated donkey anti-rabbit antibody (Jackson Immunoresearch, West Grove, PA). Cover slips were mounted using one drop of Slow Fade Gold (Molecular Probes, Eugene, OR) and edges sealed using nail polish.

Staining was visualized and images were captured using fluorescent confocal microscopy using a Nikon EFD-3 microscope (Nikon, Melville, NY) containing a Biorad MRC 1024 confocal unit utilizing Biorad LaserSharp 2000 version 5.2 software (BioRad Laboratories, Inc., Hercules, CA). Fluorescein and propidium iodide were detected using 520 nm emission/ 495 nm excitation and 617 nm emission/ 535 nm excitation, respectively. Using ImageJ software (NIH, Bethesda, MD), apoptotic cells were counted and compared to total number of cells in the image.

## Results

Following optic nerve crush, to ascertain the effects of TUDCA on retinal ganglion cell survival, Fluorogold was introduced to the damaged optic nerve via self-dissolving gel foam applied directly at the site of injury. Fluorogold is transported only in intact axons, thus, the presence fluorescent cells indicates viable RGCs. This can be seen in Fig. 1, where RGCs and microglia are fluorescing with characteristic shapes (indicated in figure). The number of fluorescently-labeled RGCs of TUDCA-treated mice were 34%, 214%, and 116% greater than vehicle treated mice at days 5, 7, and 9 post optic nerve crush respectively (Fig. 2). At day 10, 53% of RGCs were apoptotic in vehicle treated retinae; while TUDCA treated retinae were 8% apoptotic (Fig. 2).

**Figure 1.**
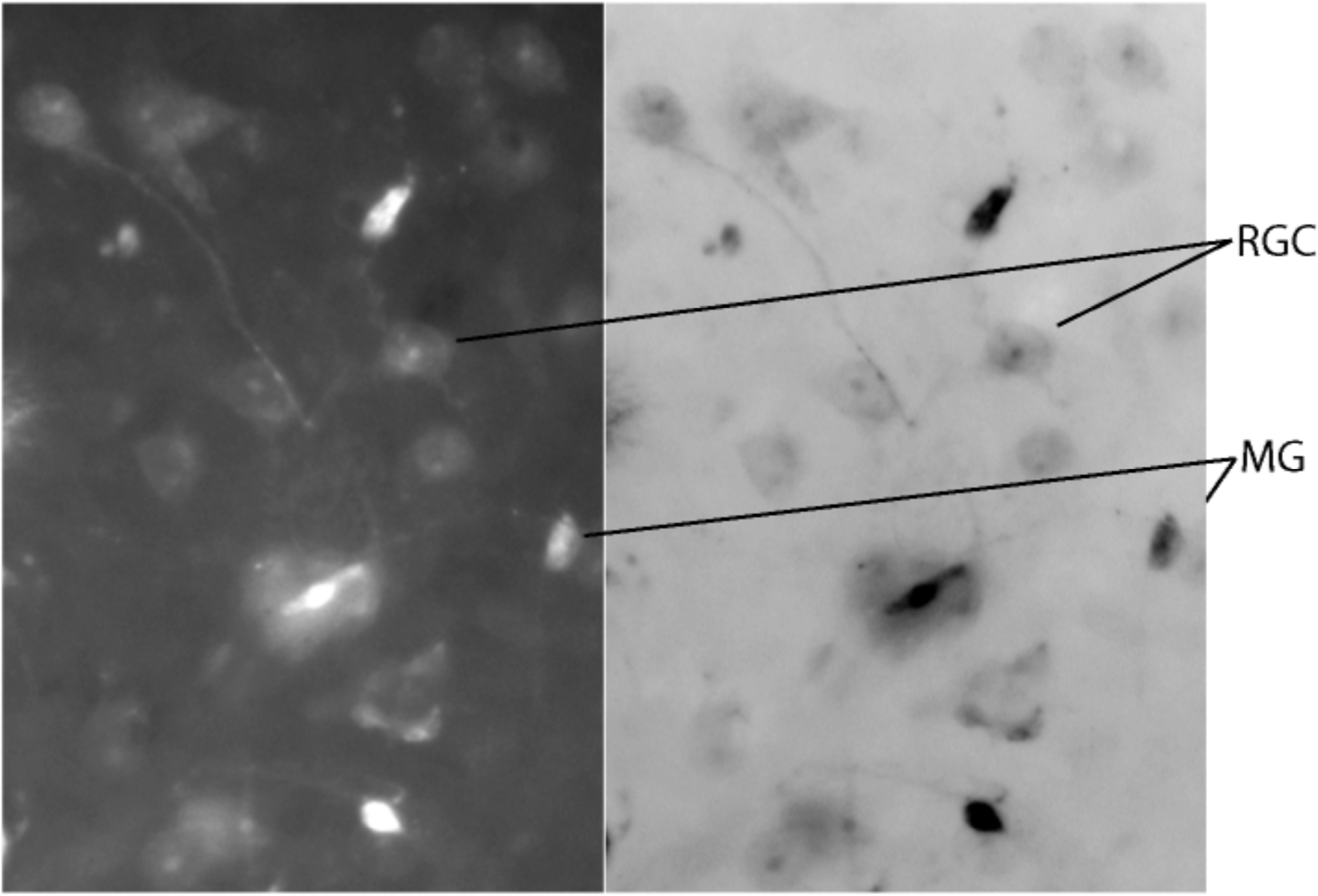
Sample photomicrograph of retina flatmount (7 days post-surgery) Representative photomicrographs of retinal flatmounts following ONC surgery. Characteristic shapes of RGC cell bodies of RGCs that have taken up Fluorogold and microglia (MG) fluorescence likely due to having ingested dead RGCs.

**Figure 2.**
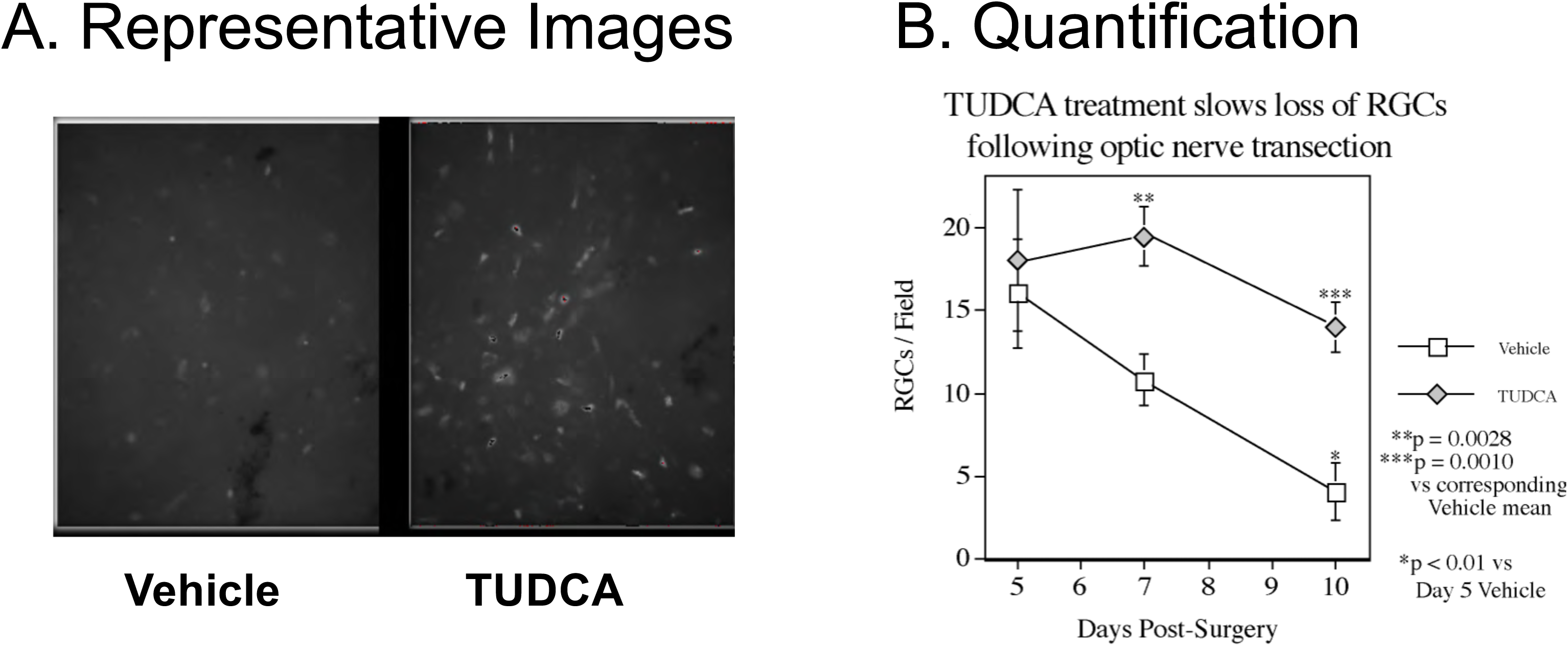
Effect of TUDCA treatment on RGC survival. A: Representative photomicrographs of retinal flatmounts following ONC surgery from mice treated with vehicle or TUDCA. Quantification shows that TUDCA treatment significantly delayed or prevented RGC loss. Statistical testing and outcomes as indicated in figure.

As shown in Fig. 3, ONC surgery resulted in TUNEL signal in all three retinal nuclear layers in some sections, whereas in other sections, TUNEL signal is largely restricted to the RGC layer. Treatment with TUDCA suppressed appearance of TUNEL all layers (Fig. 3). TUNEL-positive cells in the RGC layer were counted across entire retina sections. Sections following ONC had ’70% TUNEL-positive cells (Fig. 4). In sections from TUDCA-treated mice, only ’20% of the cells were TUNEL-positive, a statistically significant difference (Fig. 4).

**Figure 3.**
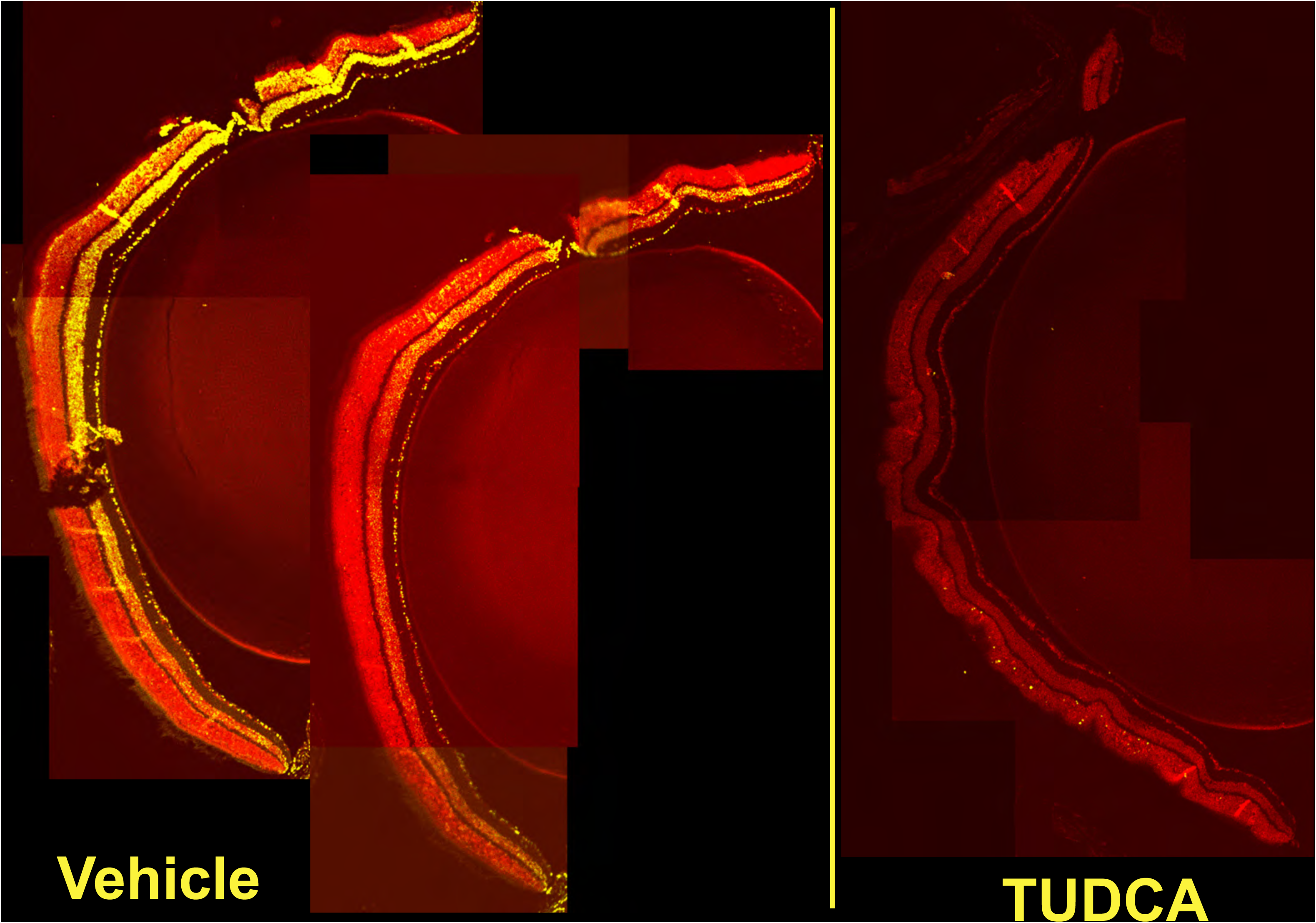
Effect of TUDCA on RGC apoptosis. Representative photomicrographs show TUNEL signal in all three retinal nuclear layers. This is prevented in all layers by TUDCA treatment.

**Figure 4.**
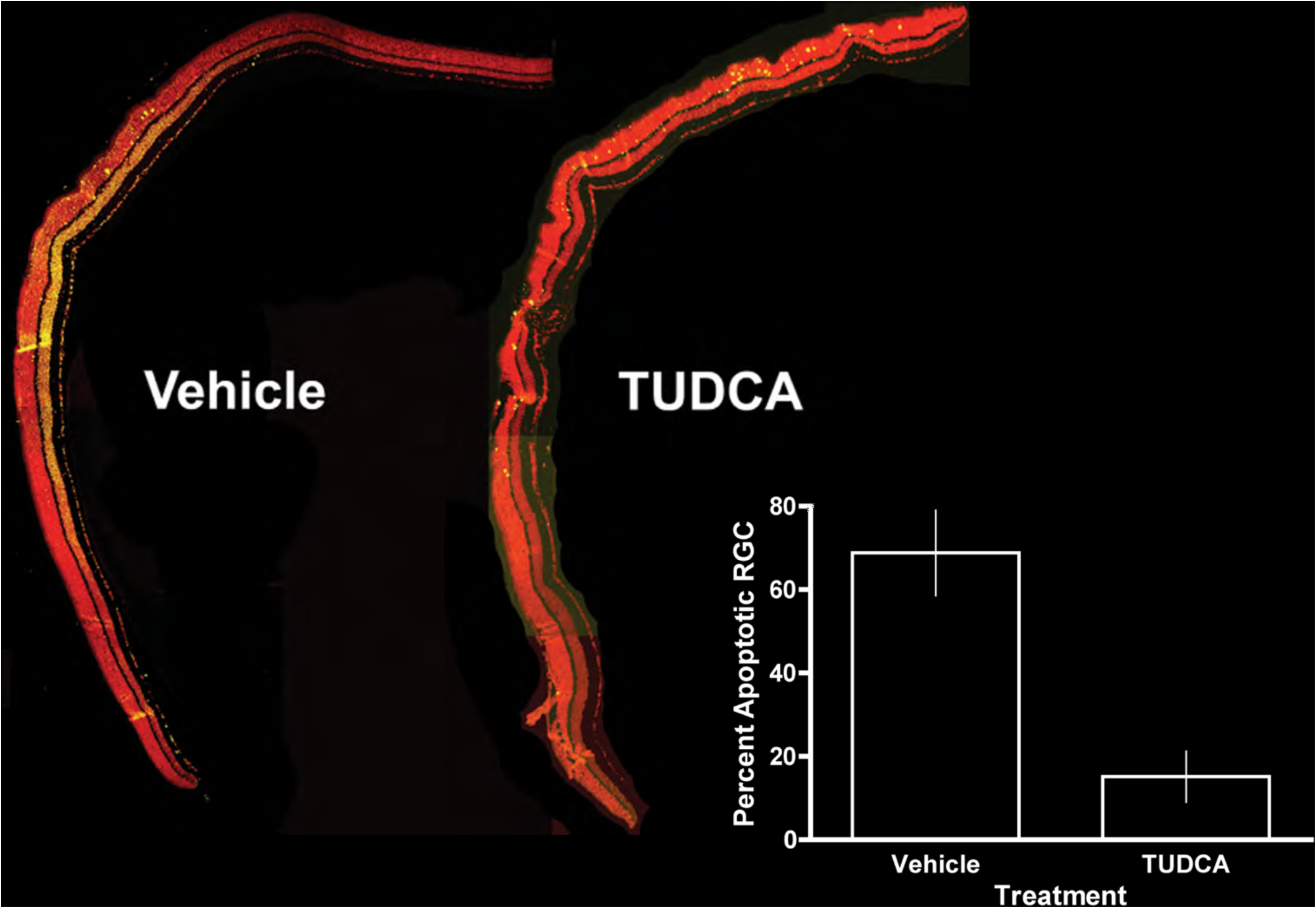
Effect of TUDCA on RGC apoptosis. TUNEL positive cells in the RGC layer were counted across entire retina sections. Sections following ONC had ’70% TUNEL-positive cells. In sections from TUDCA-treated mice, only ’20% of the cells were TUNEL-positive, a statistically significant difference (p < 0.05, Students t-test, N=6 mice/group).

In other experiments, one week after ONC, in flatmounts stained for Brn3a, immunofluorescencing cell counts were 100±4.5 cells/per field in the PBS-injected control group and 177±19 cells/per field in the TUDCA-injected group (P<0.05) (Fig. 5). Two weeks after ONC, there also were more cells labeled by RGC pan-marker RBPMS in the TUDCA-injected group (158±12 cells/per field) than in the PBS-injected control group (97±1.8 cells/per field) (P<0.05)(Fig. 6). In similar experiments, three days after ONC, in flatmounts stained for Brn3a, treatment with HIOC partially but significantly diminished ONC-induced RGC loss (Fig. 7).

**Figure 5.**
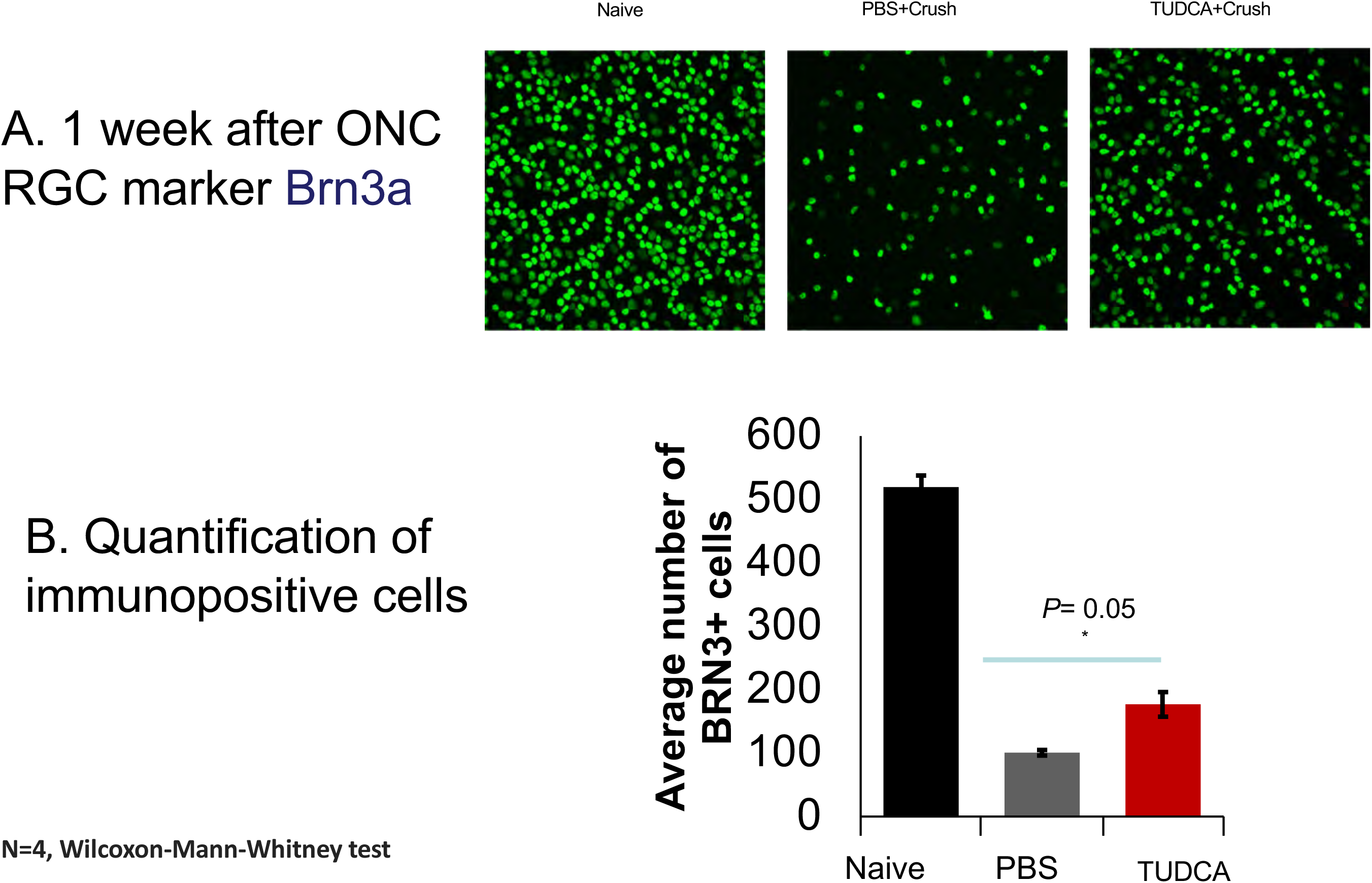
Effect of TUDCA on Brn3a-immunopositive cells. A: Representative photomicrograph fields of retinal flatmounts from naïve mice or from mice that had been treated with vehicle (PBS) or TUDCA and then undergone ONC surgery. B: Quantification shows that TUDCA treatment significantly delayed or prevented RGC loss. Statistical testing and outcomes as indicated in figure.

**Figure 6.**
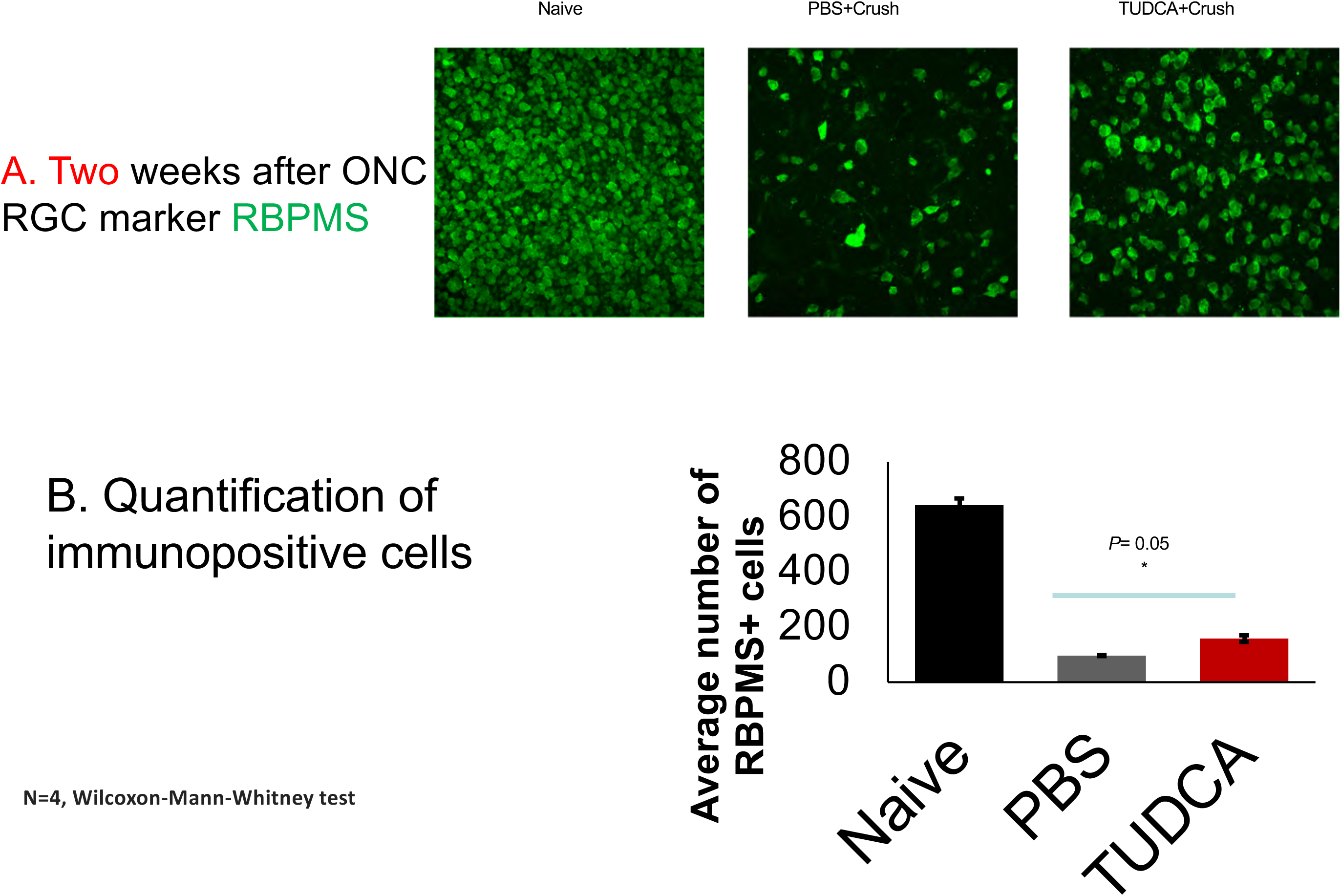
Effect of TUDCA on RBPMS-immunopositive cells. A: Representative photomicrograph fields of retinal flatmounts from naïve mice or from mice that had been treated with vehicle (PBS) or TUDCA and then undergone ONC surgery. B: Quantification shows that TUDCA treatment significantly delayed or prevented RGC loss. Statistical testing and outcomes as indicated in figure.

**Figure 7.**
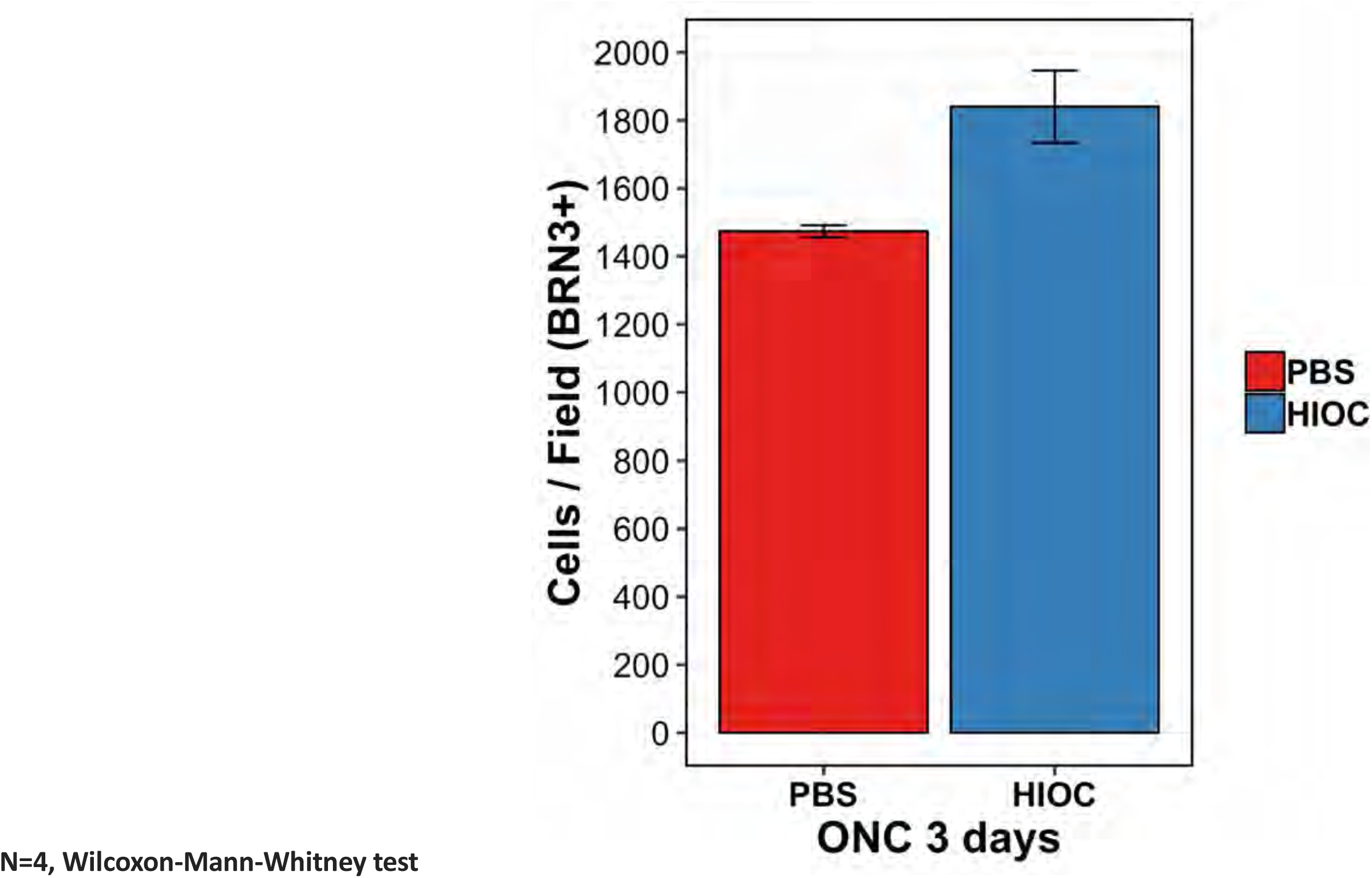
Effect of HIOC on Brn3a-immunopositive cells. HIOC treatment resulted in significantly more RGC surviving following ONC compared PBS treatment. Statistical testing and outcomes as indicated in figure. (p < 0.05, Students t-test, N=4 mice/group).

## Discussion

Systemic TUDCA treatment significantly enhanced RGC survival and suppressed apoptosis following optic nerve crush in mice. In glaucoma, retinal ganglion cells die in at least three ways: mechanical damage to the axon, vascular insufficiency, and exocytotoxic damage [22]. The surgical crush mimics the mechanical injury to retinal ganglion cell axons exiting the eye. The indirect partial occlusion of the central retinal vein, which travels in the optic nerve bundle, mimics glaucomatous vascular insufficiency. The exocytotoxic injury found in glaucoma is also mimicked in the optic nerve crush model, as the crush surgery elevates both intraocular glutamate and aspartate level [23].

The TUNEL signal in all three retinal nuclear layers of some retina sections (Figs. 3 and 4) suggests that the crush surgeries are causing non-mechanical damage, whereas the images from other crush surgeries showing TUNEL only in the RGC layer suggest that in some instances, only mechanical injury occurred. Delivery of TUDCA by i.p. injection increased survival of RGCs that are immunopositive for Brn3a (Fig. 5) or RPBMS (Fig. 6) after ONC. These data suggest that it is worthwhile to further explore protective effects of TUDCA on RGC subtypes with regards to structure and function and in additional disease models that involve RGC loss.

## Support

VA I01RX002806 (JHB); VA I21RX001924 (JHB); VARR&D C9246C (Atlanta VAMC); Abraham J and Phyllis Katz Foundation (JHB); The joint training program between Emory University School of Medicine and Xiangya School of Medicine, Central South University. China Scholarship Council (XZ); NIH R01EY028859 (JHB); NIH R01EY021592 (JMN); NIH R01EY028450 (JMN); NIH P30EY06360 (Emory); Research to Prevent Blindness (Emory).

